# Machine learning leveraging genomes from metagenomes identifies influential antibiotic resistance genes in the infant gut microbiome

**DOI:** 10.1101/185348

**Authors:** Sumayah F. Rahman, Matthew R. Olm, Michael J. Morowitz, Jillian F. Banfield

## Abstract

Antibiotic resistance in pathogens is extensively studied, yet little is known about how antibiotic resistance genes of typical gut bacteria influence microbiome dynamics. Here, we leverage genomes from metagenomes to investigate how genes of the premature infant gut resistome correspond to the ability of bacteria to survive under certain environmental and clinical conditions. We find that formula feeding impacts the resistome. Random forest models corroborated by statistical tests revealed that the gut resistome of formula-fed infants is enriched in class D beta-lactamase genes. Interestingly, *Clostridium difficile* strains harboring this gene are at higher abundance in formula-fed infants compared to *C. difficile* lacking this gene. Organisms with genes for major facilitator superfamily drug efflux pumps have faster replication rates under all conditions, even in the absence of antibiotic therapy. Using a machine learning approach, we identified genes that are predictive of an organism’s direction of change in relative abundance after administration of vancomycin and cephalosporin antibiotics. The most accurate results were obtained by reducing annotated genomic data into five principal components classified by boosted decision trees. Among the genes involved in predicting if an organism increased in relative abundance after treatment are those that encode for subclass B2 beta-lactamases and transcriptional regulators of vancomycin resistance. This demonstrates that machine learning applied to genome-resolved metagenomics data can identify key genes for survival after antibiotics and predict how organisms in the gut microbiome will respond to antibiotic administration.

**Importance:** The process of reconstructing genomes from environmental sequence data (genome-resolved metagenomics) allows for unique insight into microbial systems. We apply this technique to investigate how the antibiotic resistance genes of bacteria affect their ability to flourish in the gut under various conditions. Our analysis reveals that strain-level selection in formula-fed infants drives enrichment of beta-lactamase genes in the gut resistome. Using genomes from metagenomes, we built a machine learning model to predict how organisms in the gut microbial community respond to perturbation by antibiotics. This may eventually have clinical and industrial applications.

## Introduction

Antibiotic use has been steadily increasing over the past several decades and is correlated with the prevalence of antibiotic resistance in bacteria^1^. Widespread antibiotic resistance, in combination with the decline in development of new antibiotics, presents a major threat to human health^2^. The gut microbiome is a reservoir for antibiotic resistance genes^3^ and may be involved in the spread of resistance genes to pathogens^4^^-^^6^. Additionally, antibiotics are often prescribed to treat infections without considering how the drug will affect the gut microbial community, which can lead to negative consequences for the human host^7^. It is therefore important to study how the antibiotic resistance genes harbored by organisms in the gut microbiome impact community dynamics.

The preterm infant gut resistome is considered a research priority because premature infants are almost universally administered antibiotics during the first week of life^8^. Early life is a critically important time for community establishment^9^, and neonatal antibiotic therapies have both transient and persistent effects on the gut microbial community. Included among the many ways that antibiotics have been shown to affect the microbiome are lower bacterial diversity^10^, enrichment of *Enterobacteriaceae*^10^,^11^, reduction of *Bifidobacterium* spp.^12^, and enrichment of antibiotic resistance genes^13^, including those that have no known activity against the particular antibiotic administered^14^. Previous studies have shown that the community composition of the infant microbiome is affected by diet, with artificial formula selecting for *Escherichia coli* and *Clostridium difficile*^15^, and breast milk selecting for certain strains of *Bifidobacterium*^16^. The effect of birth mode on the microbiome is contested, with most studies finding that it has an effect on the gut microbiome^17^^-^^19^ although some show no effect^20^,^21^. Gender^22^ and maternal antibiotics before or during birth^23^^-^^25^ also influence microbiome assembly.

Here we use genome-resolved metagenomics coupled with statistical and machine learning approaches to investigate the gut resistome of 107 longitudinally sampled premature infants. We show that certain antibiotic resistance genes in particular genomes affect how clinical factors influence the gut microbiome and, in turn, how the antibiotic resistance capabilities of a gut organism influence its growth and relative abundance.

## Results and Discussion

### Antibiotic resistance of the premature infant microbiome

107 premature infants were longitudinally sampled during the first three months of life, resulting in a total of 902 samples that were sequenced and analyzed. All 107 infants received gentamicin and ampicillin during the first week of life, and 36 of those infants received additional antibiotics in the later weeks due to disease (**Table 1**). All samples were subject to Illumina short-read shotgun sequencing and the sequence data assembled using idba-ud (see methods for details). Resfams^26^ annotations of predicted amino acid sequences from the resulting scaffolds revealed that the most abundant resistance mechanisms were resistance-nodulation-division (RND) efflux pumps and ATP-binding-cassette (ABC) transporters (**Fig 1**). A slight, yet significant, decreasing trend in total antibiotic resistance potential is observed over time (*p* < 0.005). During the first week of life, empiric antibiotic therapy perturbs the microbiome though selection for antibiotic resistant organisms. This is consistent with prior results showing temporarily elevated resistance gene levels after administration of antibiotics^17^. Microbial community recovery begins following this period.

**Table 1.** All 107 infants in this analysis were in the neonatal intensive care unit (NICU) of the Magee-Women’s Hospital in Pittsburgh, PA. The infants in the received post-week antibiotics column (right) were administered antibiotics while in the NICU beyond the first week of life due to late-onset sepsis, necrotizing enterocolitis, or another disease. The antibiotics administered included ampicillin, gentamycin, vancomycin, cefotaxime, cefalozin, amoxicillin, clindamycin, naficillin, ofloxacin, and piperacillin-tazobactam.

|  | No antibiotics after the first week | Received post-week antibiotics |
| --- | --- | --- |
| Samples | 604 | 298 |
| Infants | 71 | 36 |
| Received breast milk | 52 | 32 |
| Delivered by C-section | 54 | 22 |
| Male sex | 34 | 17 |
| Maternal antibiotics | 24 | 20 |

**Figure 1.**
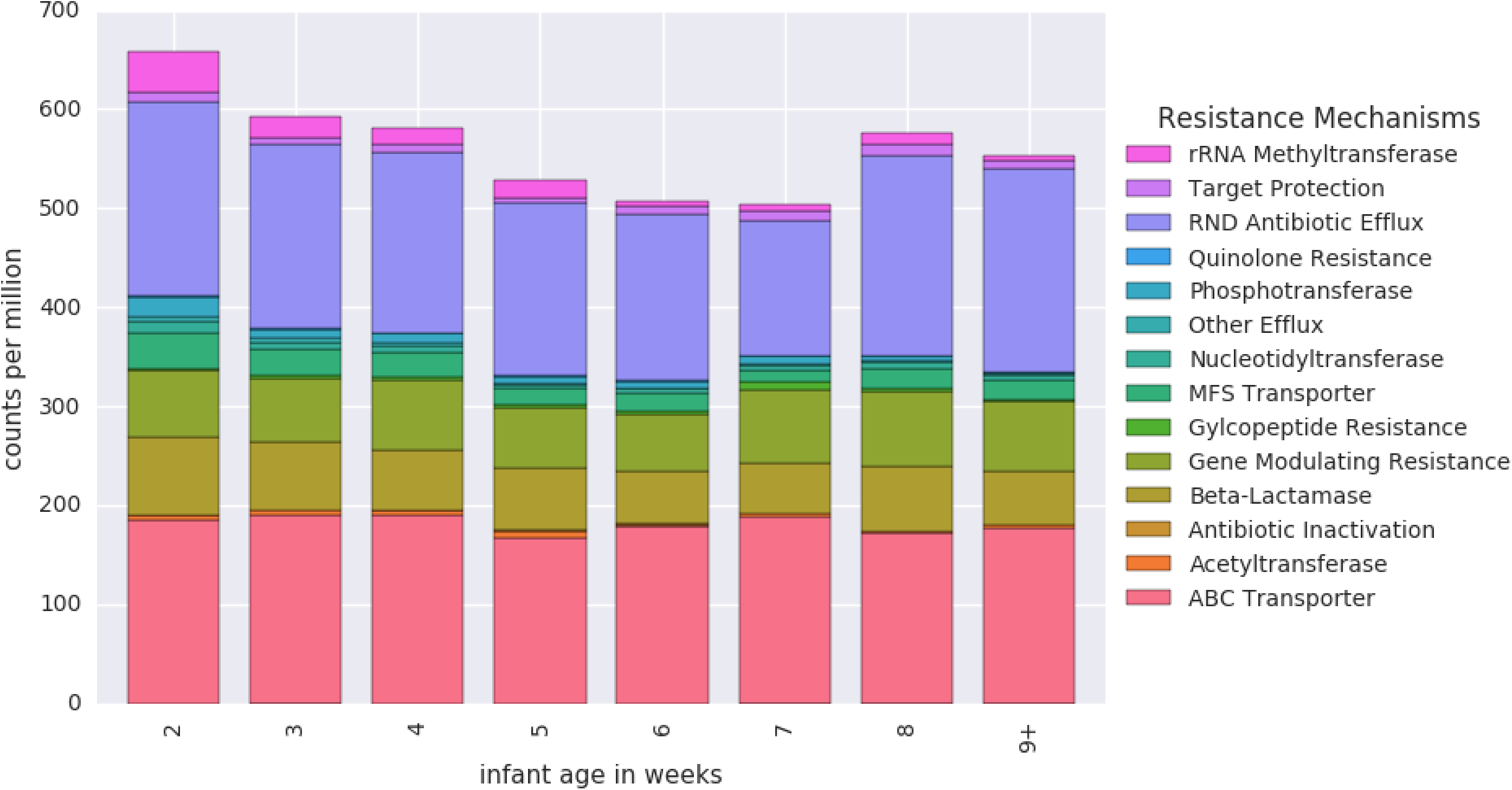
Antibiotic resistance genes of the premature infant gut microbial community. Genes were annotated through hidden Markov model searches of the Resfams database and categorized based on resistance mechanism. During the time period studied, the total resistance content of the premature infant gut microbiome has a slight negative correlation with age (Pearson’s *r* = −0.1, p = 0.003.)

### Formula feeding influences the gut resistome through strain-level selection

Permutational multivariate analysis of variance (PERMANOVA) tests, which discern and isolate the effects of factors through partitioning of variance^27^, were performed to investigate the effect of feeding regimen, delivery mode, gender, maternal antibiotics, and the infant’s current antibiotic therapy on the resistome. Tests were performed on the resistomes of samples taken at weeks two, four, and six (see methods for details). At week two, formula-fed infants did not have a significantly different distribution of antibiotic resistance genes compared to infants that received breast milk. However, a difference was detected at weeks four and six (*p* < 0.05), accompanied by an increase in effect size as assessed by PERMANOVA F-statistic (**Table S1**). This signals divergence of the resistomes of formula-fed and breast-fed infants over time. Interestingly, other factors such as birth mode and antibiotic administration were not associated with a significant difference in resistance gene distribution.

Random forest models were used to classify resistomes as either belonging to a formula-fed baby or a breast-fed baby, and we used the trained model’s feature importance scores to select resistance genes for further study (Table 2). One out of the four selected resistance genes was significantly associated with a feeding group: Class D beta-lactamase was enriched in formula-fed infants (*p* < 0.05) (Fig 2A). The mexX gene, which encodes for an efflux pump subunit, exhibited a comparatively higher feature importance score, but there was no significant difference in its abundance between feeding groups (Table 2). This suggests that the oft-used practice of utilizing feature importance scores of random forests or other machine learning ensemble methods to identify markers of differences between groups^28^^-^^30^, although potentially informative, may not necessarily rank features in the order that most meaningfully reflects their contributions. This is likely due to a preference for correlated predictor variables^31^.

**Figure 2.**
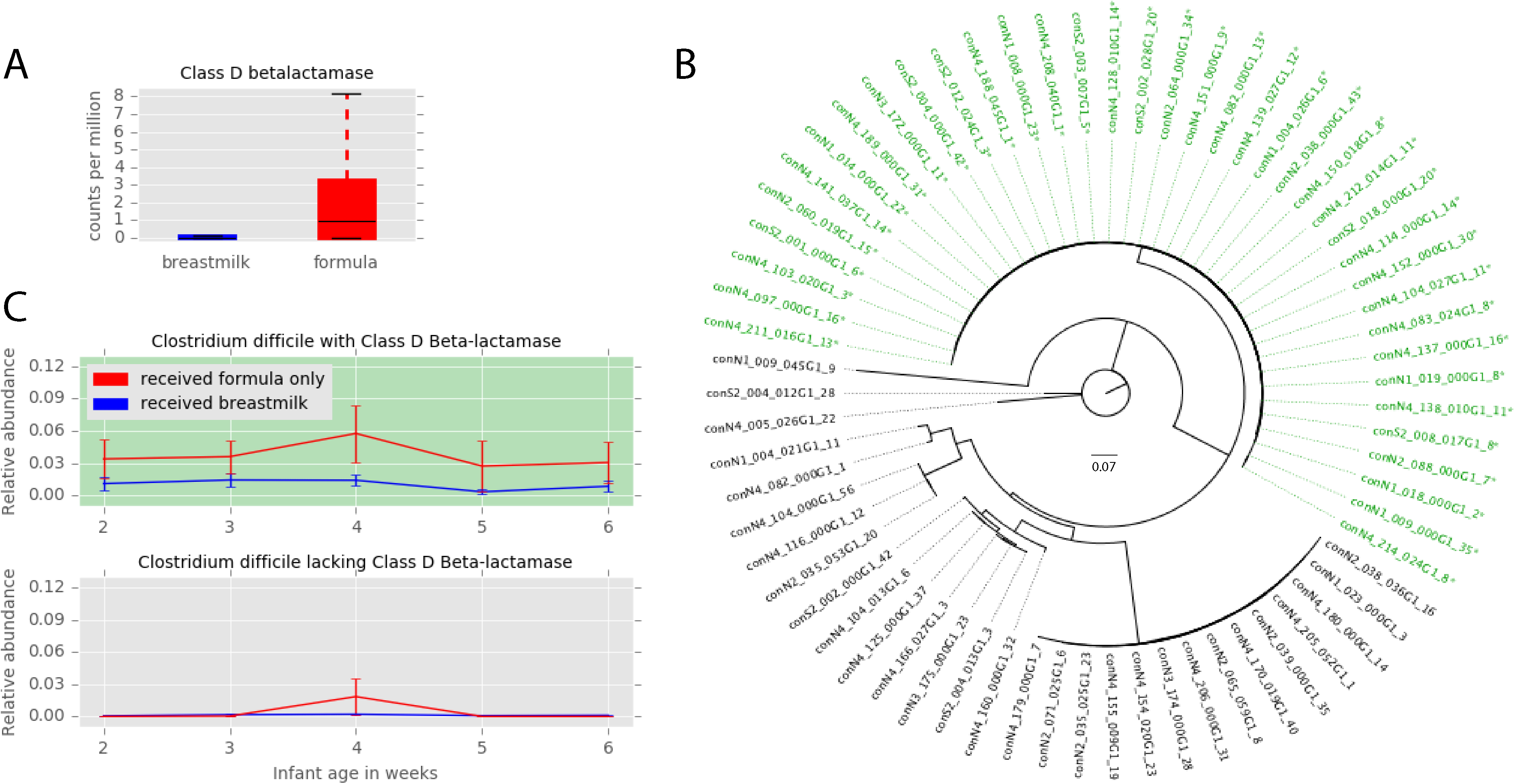
Formula feeding affects the resistome. (**A**) Class D beta-lactamase is enriched in formula fed infants at four weeks of age (Mann-Whitney U = 66, Bonferonni corrected *p* = 0.031). (**B**) Phylogenetic tree of *Clostridium difficile* genomes based on the ribosomal protein S3 gene. Names of genomes harboring a Class D beta-lactamase are colored green and labeled with an asterisk. (**C**) The relative abundance of *C. difficile* genomes with Class D beta-lactamase in formula-fed and breast-fed infants (top, *n* = 38) and the relative abundance of *C. difficile* genomes lacking Class-D betalactamase in formula-fed and breast-fed infants (bottom, *n* = 29).

**Table 2.**
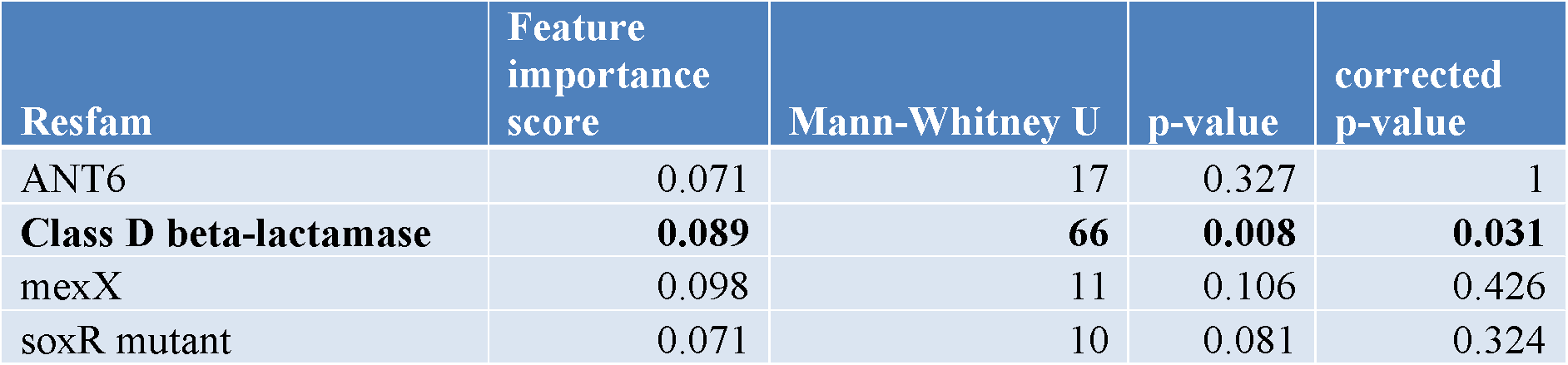
Mann-Whitney U tests were performed on features selected by the random forest model gini importance metric after training on resistomes of formula-fed infants and breast-fed infants. Bonferonni corrections were applied to the p-values to adjust for multiple testing.

Genome-resolved analysis revealed that Class D beta-lactamase genes are most frequently carried by *Clostridium difficile*. Of the 67 *C. difficile* genomes in the dereplicated dataset, 38 of these organisms harbor a Class D beta-lactamase gene. Phylogenetic analysis reveals that these 38 organisms are very closely related (Fig 2B). To ascertain if this *C. difficile* strain is involved in the enrichment of Class D beta-lactamase in the formula-fed infant gut resistome, the relative abundance of *C. difficile* with and without a Class D beta-lactamase gene in the gut microbiome of breast-fed and formula-fed infants was assessed. In infants that only receive formula, *C. difficile* with Class D beta-lactamase is consistently more abundant than *C. difficile* lacking this gene; while in infants that receive breast milk, both types of *C. difficile* are low in relative abundance (Fig 2C). Prior studies have reported an increased abundance of *C. difficile* in the gut microbiomes of formula-fed infants^15^, but here we reveal that formula feeding selects for a particular *C. difficile* strain.

Class D beta-lactamase hydrolyzes beta-lactam antibiotics^32^, and there is no known connection between host diet and its antibiotic resistance function. Since it is thus unlikely that Class D beta-lactamase offers a selective advantage to organisms in the gut of formula-fed infants, other possibilities were explored. Pairwise correlations of the Resfams and KEGG metabolism modules in *C. difficile* genomes revealed that one KEGG module, the cytidine 5′-monophospho-3-deoxy-D-*manno*-2-octulosonic acid (CMP-KDO) biosynthesis module, was perfectly correlated with the presence of the Class D beta-lactamase gene. CMP-KDO catalyzes a key reaction in lipopolysaccharide biosynthesis^33^. Further inspection of the KEGG annotations revealed that only one gene from this module was present in *C. difficile*: arabinose-5-phosphate isomerase. This gene typically occurs in Gram-negative bacteria, where it plays a role in synthesis of lipopolysaccharide for the outer membrane^34^, yet a recent study identified arabinose-5-phosphate isomerase in a Gram-positive organism, *Clostridium tetani*^35^. Although the function of this gene in Gram-positive bacteria is unknown, it is hypothesized to be a regulator and may modulate carbohydrate transport and metabolism^35^. If so, *C. difficile* (Gram-positive) strains with arabinose-5-phosphate isomerase may have a competitive advantage because they are able to rapidly respond to availability of the carbohydrates that are abundant in formula.

### Major facilitator superfamily pumps are associated with increased replication

A previous analysis revealed that antibiotic administration is associated with elevated replication rates of gut organisms, which was hypothesized to be due to high resource availability after elimination of antibiotic susceptible strains^36^. We show here that a sample’s mean replication index in the days following antibiotic treatment is positively correlated with total resistance gene content (*p* < 0.05) (Fig 3A). The replication index, or iRep value^36^, is influenced by the fraction of cells in the population that are replicating. Our results suggest that the elevated iRep values after antibiotic treatment are likely not solely explained by resource availability, but rather appear to be influenced by the organisms’ resistance gene sets.

**Figure 3.**
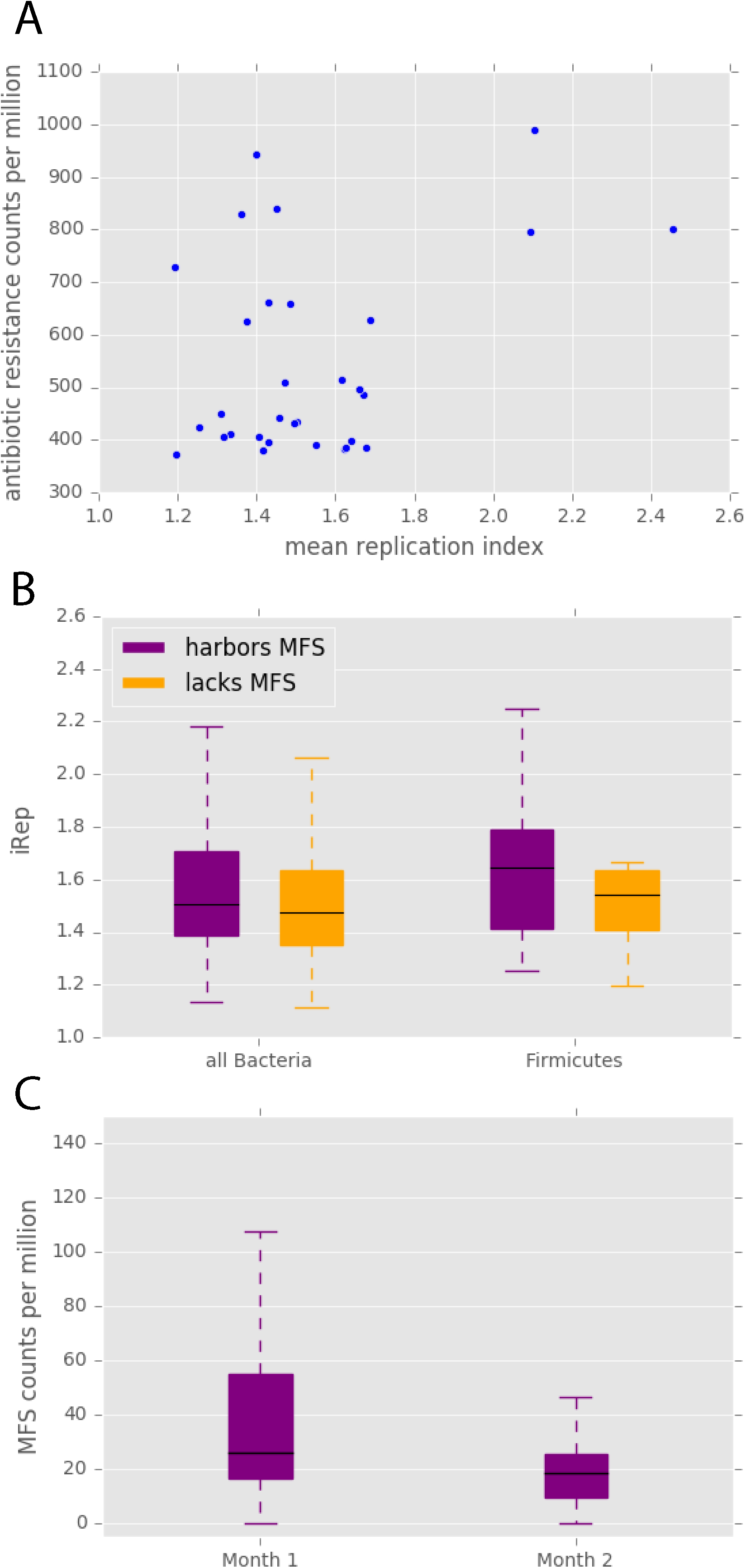
Antibiotic resistance and replication. (**A**) In samples taken within five days after antibiotic treatment, the antibiotic resistance potential of that sample is correlated with its mean replication index (Pearson’s *r* = 0.39, *p* = 0.03). (**B**) In infants that did not receive antibiotics after the first week of life, bacteria harboring at least one major facilitator superfamily (MFS) transporter gene have significantly higher iRep values (Mann-Whitney U= 827176.0, *p* = 1.55 × 10^−5^), and this pattern is apparent within the Firmicutes phylum (Mann-Whitney U = 136756.0, *p* = 0.0002). (**C**) MFS transporters are enriched in the first month of life compared to the second month of life (Wilcoxon signed-rank W = 165, *p* = 0.02). All p-values are Bonferonni corrected.

To characterize the effect of antibiotic resistance genes on iRep values in isolation from the confounding effects of antibiotics, we studied infants that did not receive any antibiotics after the first week of life. In these infants, organisms carrying genes for major facilitator superfamily (MFS) transporters have significantly higher iRep values than those that do not have MFS genes *(p* < 5 x 10^−5^) (Fig 3B). As there are known differences in median iRep values among phyla^36^, the comparison was repeated within each phylum that contained members with and without MFS genes. The trend of higher iRep values for organisms with MFS was most apparent in Firmicutes (*p* < 5 x 10^−4^) (Fig 3B). Therefore, the presence of these antibiotic resistance genes appears to inherently increase replication, even when no antibiotics are being administered. This could be due to protection from antibiotics being produced at a low level by other gut organisms^37^ or a result of MFS pumps’ naturally beneficial physiological functions^38^. We also acknowledge that this finding may simply reflect high incidence of organisms with MFS genes during periods of fast replication without a causal link.

We discovered that MFS gene levels in the resistome are significantly lower in the second month of life compared to the first (*p* < 0.05) (Fig 3C). MFS is the only resistance mechanism category for which this is true. The comparative enrichment of MFS genes in early in life may be a response to the empiric antibiotics (ampicillin and gentamicin) administered to all premature infants during the first week. The decline in MFS genes supports the established phenomenon of an eventual decrease of antibiotic resistance gene levels in the microbiome after the cessation of antibiotic therapy^17^.

### A model that predicts an organism’s response to vancomycin and cephalosporins

We modeled the relationship between gene content of a gut organism and its direction of change in relative abundance (increase vs. decrease) after a premature infant is administered a combination of glycopeptide (vancomycin) and beta-lactam (cephalosporin, either cefotaxime or cefazolin) antibiotics. Principal component analysis was performed on Resfams^26^ and KEGG^39^ annotations to generate a low-dimensional representation of each organism’s metabolic potential and resistance potential. The first five principal components (PCs) cumulatively explained 48% of the variation in the dataset. Using these PCs as input, the AdaBoost-SAMME algorithm^40^ was applied, with decision tree classifiers as base estimators. The model, trained on 70% of the data, performed extremely well on the validation set, with a precision of 1.0 and recall of 1.0, indicating that every genome was correctly classified. Because the validation set was utilized for testing during the preliminary stages of model development, the model was also evaluated with a final test set, on which it achieved 0.9 precision and 0.7 recall.

Of the features that acted as the strongest contributors to each of the PCs, five genes with a tendency to occur in microbes that increase in relative abundance after antibiotic treatment were identified (Fig 4). One of these is subclass B2 beta-lactamase. Subclass B2 beta-lactamase of *Aeromonas* spp. has been shown to hydrolyze carbapenems and displays much lower levels of resistance to cephalosporins^41^. Due to its substrate specificity for carbapenems, it was surprising that this beta-lactamase was among the top predictors of an organism’s ability to persist after treatment with cephalosporins. However, the substrate specificity of an antibiotic resistance gene can depend on the context of that gene^42^, and a single base substitution in a beta-lactamase gene can alter substrate specificity^43^. Our results imply that in some gut organisms, beta-lactamases falling into the B2 subclass may confer resistance to cephalosporins.

**Figure 4.**
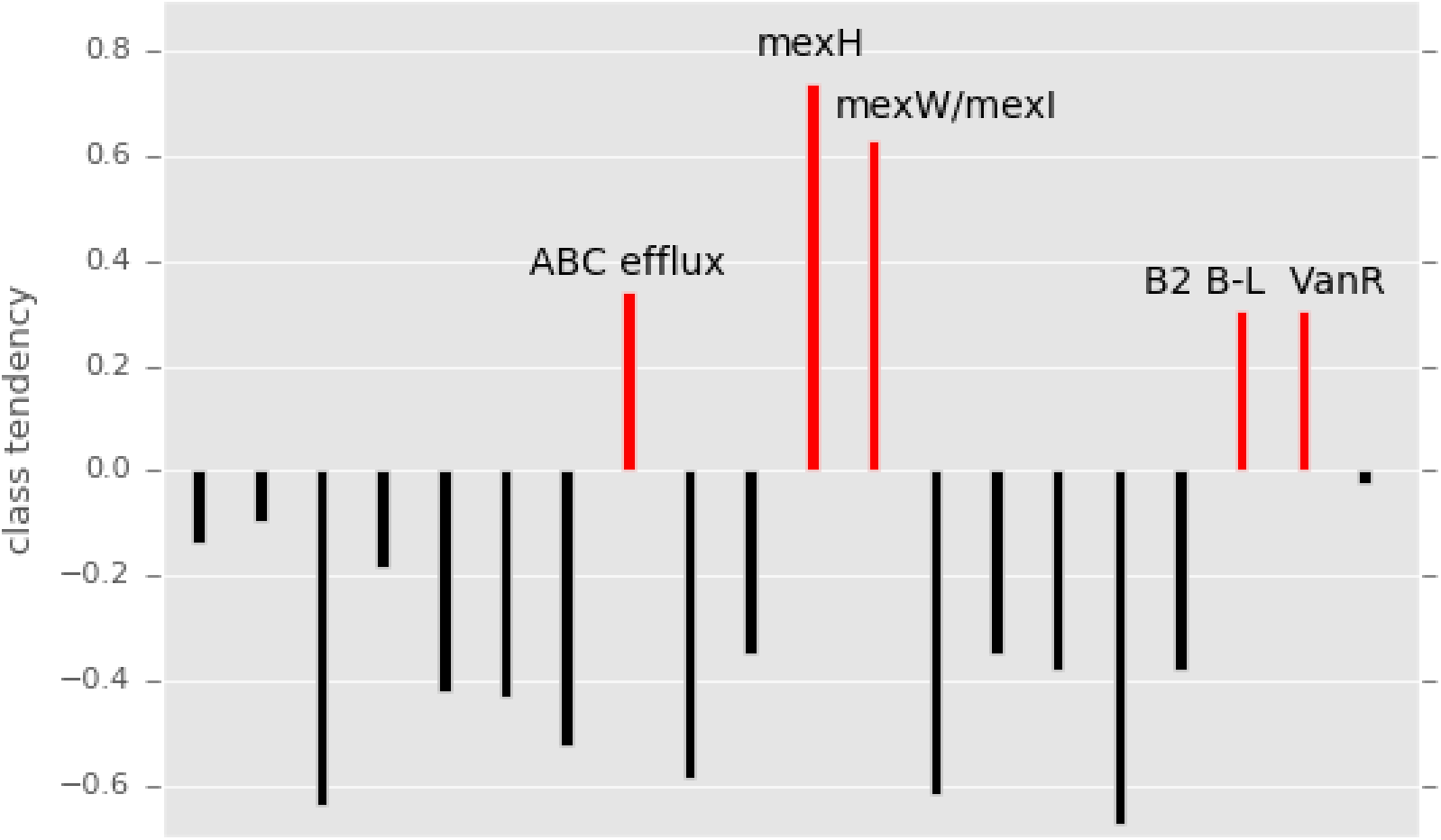
The tendency for genes to occur in the class of genomes that increased in relative abundance after antibiotics. Genes and modules strongly contributing to the principal components used in the machine learning model were identified, and class tendency was calculated using the ratio of the gene’s prevalence in the increase group to its prevalence in the decrease group. Genes associated with the increase class of genomes are colored red.

Furthermore, our model shows that a gene linked to vancomycin resistance, vanR, is among the genes predictive of an organism’s propensity to increase in relative abundance after antibiotic treatment (Fig 4). VanR is the transcriptional activator of an operon encoding genes involved in peptidoglycan modification (VanH, VanA, and VanX)^44^. This gene cluster confers resistance because peptidoglycan is the target of vancomycin, and the antibiotic cannot bind to the modified version. VanR is essential for the initiation of the vancomycin resistance operon promoter^45^, which may explain why this gene was so crucial in our model.

In addition to genes specifically encoding for resistance to beta-lactams or glycopeptides, efflux pumps and transporters were also strong contributors to the PCs used as input to the model. Mex genes (of the resistance nodulation cell division family of drug efflux pumps) and ATP-binding cassette (ABC) transporter genes were associated with microbes that increase in relative abundance after antibiotics (Fig 4). Multidrug efflux pumps are essential for the intrinsic drug resistance of many bacteria, and overexpression of the genes for these pumps leads to elevated resistance levels^46^. Among possible directions for future research is inclusion of transcriptomics data to develop a model with even more robust predictive capability.

Previous studies have utilized data from 16S rRNA gene amplicon sequencing or read-based metagenomics of the human microbiome to predict life events of the human host using machine learning or other modeling techniques^47^,^48^. However, read-based metagenomics lacks resolution at the genomic level, and due to strain-level differences in antibiotic resistance^48^, taxonomy data from marker gene studies cannot be used to predict how particular organisms in a community will respond to antibiotics. Here, for the first time, we utilize the high-dimensional data obtained by reconstructing genomes from metagenomes to make predictions about the future states of individual gut microbes. This has tremendous potential for application in the fields of medicine and microbial ecology. For example, such a model can be used before administering drugs to a patient to verify that a particular combination of antibiotics will not lead to overgrowth of an undesirable microbe. In other environmental systems, various types of perturbations can be modeled to make predictions about how they will influence key members of microbial communities. The trained models can then be used to inform decisions in areas like agriculture and contaminant removal, where individual organisms may be of great importance^51^,^52^. Our study serves as a proof of concept for this application of machine learning used in conjunction with genome-resolved metagenomics to derive biological insight.

## Materials and Methods

### Sample collection, sequencing, assembly, and gene prediction

Fecal samples were collected from 107 infants of the NICU at the Magee Women’s hospital during the first three months of life. Briefly, DNA was extracted using the PowerSoil DNA isolation kit (MoBio Laboratories, Carlsbad, CA, USA) and sequenced using the Illumina HiSeq platform. Details on sample recovery, extraction, library preparation, and sequencing have been previously reported ^53^^-^^55^. Using default parameters for all the programs, the reads were trimmed with Sickle (https://github.com/najoshi/sickle), cleared of human contamination following mapping to the human genome with Bowtie2^56^, and assembled with idba_ud^57^. Additionally, idba_ud was used to generate co-assemblies for each infant by simultaneously assembling all the samples belonging to the infant. Prodigal^58^ run in the metagenomic mode was used for gene prediction.

### Genome recovery and relative abundance calculation

For each infant, reads from all samples from that infant were mapped to all individual assemblies from that infant as well as the infant’s co-assembly using SNAP^59^.

Coverage of scaffolds was calculated and used to run concoct^60^ with default parameters on all individual assemblies and co-assemblies. To remove redundant bins, all bins recovered from each infant were dereplicated using dRep^61^ v0.4.0 with the command: *dRep dereplicate_wf–S_algorithm gANI-comp 50-con 25-str 25-l 50000-pa .9-nc .1*.

Using Bowtie2^56^, the reads from each sample were mapped to the set of genomes that were recovered from that particular infant. The read mapping output files were used to calculate the average coverage of each genome in each sample, and the coverage values were converted to relative abundance by utilizing the read length, total number of reads in the sample, and genome length.

### iRep calculation

For each sample, a set of representative genomes was first chosen from the complete collection of dereplicated genomes. First, all genomes were clustered at 98% ANI using dRep^61^. A pangenome was then generated for each of these clusters using PanSeq^62^, creating a list of fragments representing the entire sequence-space of each cluster. All pangenomes of all clusters were merged, and reads from all samples were mapped to the resulting pangenome set using SNAP^59^. By analyzing the coverage of all fragments in the pangenome set, the breadth of each genome in each sample was calculated (number of genome fragments > 1x coverage / total genome fragments). Genomes with less than 85% breadth were removed from analysis. For all remaining genomes, the genome from each cluster with the highest breadth was added to that sample’s representative genome list.

Next, reads from each sample were mapped to its representative genome list using bowtie2^56^ default parameters. iRep^36^ was run on the resulting mapping files using default parameters and without GC correction. Only values that passed iRep’s default filtering and were < 3 were considered for analysis.

### Annotation

The amino acid sequences of genes predicted by the metaProdigal gene finding algorithm^58^ were searched against Resfams^26^, an antibiotic resistance gene specific profile hidden Markov model (HMM) database using the *hmmscan* function of HMMER v 3.1b2^63^. The *–cut_ga* option was used to set the reporting and inclusion limits as the profile-specific gathering threshold, which have been manually optimized on a profile-by-profile basis to ensure Resfams prediction accuracy^26^. The Resfams annotation output and the coverage of each scaffold that had a hit to a Resfams profile were used to generate sample resistance gene summaries. Each sample resistance gene summary, which represents the antibiotic resistance potential of a particular infant gut microbiome at a particular point in time, contains the counts per million (CPM) for each of the 170 antibiotic resistance gene families in the Resfams database. Additionally, genome resistance gene profiles that indicated the count of each resistance gene were developed for each genome.

To gather general metabolism data, all binned sequences were searched against Kyoto Encyclopedia of Genes and Genomes (KEGG)^39^ HMMs and the results were parsed for genome profiling. This resulted in a KEGG metabolism profile for each organism that displayed the fraction of each KEGG module encoded by that genome.

### Statistical and computational analysis

To evaluate the effect of feeding regimen, delivery mode, gender, maternal antibiotics, and the infant’s current antibiotic therapy, three cross-sectional PERMANOVA^64^ tests for weeks two, four, and six were performed using the *adonis2* function of the *vegan* package in R^65^. For each infant, the first sample of each week was identified and the resistance gene summary of that sample was included in the PERMANOVA. If antibiotics were being administered on the day of sampling, the infant was labeled as currently receiving antibiotics. Infants that were exclusively fed breast milk and infants that were given breast milk at any point were both labeled as receiving breast milk. The Bray-Curtis dissimilarity metric was used and 9,999 permutations were performed to assess the marginal effects of the terms. The factor revealed to have a significant difference in antibiotic resistance gene content (*p* < 0.05) was selected for continued analysis. To identify antibiotic resistance genes associated with either formula feeding or breast milk during the weeks indicated by the PERMANOVA results, the infant’s diet was used to classify sample resistance gene summaries using random forest models^66^. Mann-Whitney U tests were performed on Resfams that had feature importance scores above 0.07 in the random forest models, as calculated by the Gini importance metric. Genomes containing resistance genes significantly associated with a particular feeding type, along with genomes of the same species lacking these genes, were further investigated. The ribosomal protein S3 (RPS3) genes for each genome were identified using rp16.py (https://github.com/christophertbrown/bioscripts/blob/master/bm/rp16.py). The RPS3 nucleotide sequences were aligned with MUSCLE^67^ using default parameters, and a maximum-likelihood phylogenetic tree was built with RAxML^68^. Pairwise Pearson correlations of Resfams with KEGG modules within these genomes were calculated.

The Pearson correlation of mean replication index (iRep) for a sample and the sample’s total resistance gene content was determined for samples collected within five days following antibiotic treatment. The replication rates of organisms harboring antibiotic resistance genes were compared to those lacking resistance genes of the same category. The first sample from a particular infant from the first month of life was matched with the same infant’s first sample from the second month of life, and Wilcoxon signed rank tests were performed to evaluate the change in abundance of each resistance mechanism. All p-values were Bonferonni corrected for multiple testing.

Infants for which there was a sample taken both before and after post-week antibiotic treatment were identified and the before and after samples were selected. Genomes from these samples were identified and labeled as either increasing or decreasing in relative abundance from the pre-antibiotic sample to the post-antibiotic sample. Using scikit-learn^66^, development of a machine learning model to predict the direction of change in relative abundance for each genome based on its Resfams and KEGG metabolism was attempted; yet an adequate model could not be developed, presumably due to variation in the ways that organisms respond to different antibiotic combinations. Therefore, the dataset was narrowed to include the six infants that received either cefotaxime or cefazolin (both cephalosporin antibiotics) in conjunction with vancomycin. 70% of the genomes obtained from these infant samples were used for training, 15% was used as a validation set for model improvement, and 15% was held out as a final test set. Several attempts to improve model performance through algorithm choice, feature engineering, and parameter tuning were applied, and the model that exhibited the best results with regard to precision and recall was selected. This model was then used to make predictions on the final test set. Each feature constructed for the model was a principal component of the Resfams and KEGG metabolic data, and the genes/modules contributing most strongly to each of these principal components were identified. The tendency for each of the genes and modules to occur in the increase class was calculated by adding-1 to the gene’s mean value in the increase class divided by its mean value in the decrease class.

### Data availability

The dataset used is comprised of 597 previously reported samples^53^^-^^55^, as well as 305 new samples. These samples are available at NCBI under accession number SRP114966 (https://www.ncbi.nlm.nih.gov/sra/?term=SRP114966). The code for the analysis, along with all the data and metadata used in the analysis, is hosted at https://github.com/SumayahR/antibiotic-resistance.

## Acknowledgements

Funding was provided through the National Institutes of Health (NIH) under grant RAI092531A and the Alfred P. Sloan Foundation under grant APSF-2012-10-05. This work used the Vincent J. Coates Genomics Sequencing Laboratory, supported by NIH S10 OD018174 Instrumentation Grant. The study was approved by the University of Pittsburgh Institutional Review Board (IRB) (Protocol PRO12100487).

We acknowledge Robyn Baker for recruiting infants, and Brian Firek for performing DNA extractions. We would also like to thank David Burstein for the KEGG HMM annotation pipeline, and Christopher Brown for scripts to identify ribosomal proteins and to calculate genome coverage.

## Author Contributions

MJM and JFB conceived of the project. MRO processed the sequence data, including binning and de-replication of genomes. SFR performed the computational analysis and interpreted the results. SFR and JFB wrote the manuscript, and all authors contributed to manuscript revisions.

## References

1. Goossens, H., Ferech, M., Vander Stichele, R. & Elseviers, M. Outpatient antibiotic use in Europe and assaciation with resistance: a cross-national database study. Lancet 365, 579–587 (2005).

2. Spellberg, B. et al. The Epidemic of Antibiotic-Resistant Infections: A Call to Action for the Medical Community from the Infectious Diseases Society of America. Clin. Infect. Dis. 46, 155–164 (2008).

3. Penders, J., Stobberingh, E. E., Savelkoul, P. H. M. & Wolffs, P. F. G. The human microbiome as a reservoir of antimicrobial resistance. Front. Microbiol. 4, 1–7 (2013).

4. Van Braak, N. Den et al. Molecular characterization of vancomycin-resistant enterococci from hospitalized patients and poultry products in the Netherlands. J. Clin. Microbiol. 36, 1927–1932 (1998).

5. Teuber, M., Meile, L. & Schwarz, F. Acquired antibiotic resistance in lactic acid bacteria from food. Antonie van Leeuwenhoek, Int. J. Gen. Mol. Microbiol. 76, 115–137 (1999).

6. Simonsen, G. S., Lvseth, A., Dahl, K. H. & Kruse, H. Enterococci and vanA Resistance Elements between Chicken. Microb. Drug Resist. 4, 313–318 (1998).

7. Langdon, A., Crook, N. & Dantas, G. The effects of antibiotics on the microbiome throughout development and alternative approaches for therapeutic modulation. Genome Med. 8, 39 (2016).

8. Clark, R., Bloom, B., Spitzer, A. R. & Gerstmann, D. R. Reported Medication Use in the Neonatal Intensive Care Unit: Data From a Large National Data Set. Pediatrics 29, 18–20 (2014).

9. Faith, J. J. et al. The Long-Term Stability of the Human Gut Microbiota. Science (80-.). 341, 1237439 (2013).

10. Greenwood, C. et al. Early empiric antibiotic use in preterm infants is associated with lower bacterial diversity and higher relative abundance of enterobacter. J. Pediatr. 165, 23–29 (2014).

11. Arboleya, S. et al. Intestinal Microbiota Development in Preterm Neonates and Effect of Perinatal Antibiotics. J. Pediatr. 166, 538–544 (2015).

12. Tanaka, S. et al. Influence of antibiotic exposure in the early postnatal period on the development of intestinal microbiota. FEMS Immunol Med Microbiol 56, 80–87 (2009).

13. Jernberg, C., Löfmark, S., Edlund, C. & Jansson, J. K. Long-term ecological impacts of antibiotic administration on the human intestinal microbiota. ISME J. 1, 56–66 (2007).

14. Gibson, M. K. et al. Developmental dynamics of the preterm infant gut microbiota and antibiotic resistome. Nat. Microbiol. 1, 16024 (2016).

15. Penders, J. et al. Quantification of Bifidobacterium spp., Escherichia coli and Clostridium difficile in faecal samples of breast-fed and formula-fed infants by real-time PCR. FEMS Microbiol. Lett. 243, 141–147 (2005).

16. Costello, E. K., Stagaman, K., Dethlefsen, L., Bohannan, B. J. M. & Relman, D. A. The Application of Ecological Theory Toward an Understanding of the Human Microbiome. Science (80-.). 336, 1255–1263 (2012).

17. Yassour, M. et al. Natural history of the infant gut microbiome and impact of antibiotic treatment on bacterial strain diversity and stability. Sci. Transl. Med. 8, 343ra81 (2016).

18. Wampach, L. et al. Colonization and succession within the human gut microbiome by archaea, bacteria and microeukaryotes during the first year of life. Front. Microbiol. 8, 738 (2017).

19. Penders, J. et al. Factors Influencing the Composition of the Intestinal Microbiota in Early Infancy. Pediatrics 118, 511–521 (2006).

20. Stewart, C. J. et al. Cesarean or Vaginal Birth Does Not Impact the Longitudinal Development of the Gut Microbiome in a Cohort of Exclusively Preterm Infants. Front. Microbiol. 8, 1–9 (2017).

21. Chu, D. M. et al. Maturation of the infant microbiome community structure and function across multiple body sites and in relation to mode of delivery. Nat. Med. 23, 314–326 (2017).

22. Cong, X., Xu, W., Janton, S., Henderson, W. A. & Matson, A. Gut Microbiome Developmental Patterns in Early Life of Preterm Infants?: Impacts of Feeding and Gender. PLoS One 11, e0152751 (2016).

23. Mshvildadze, M. et al. Intestinal Microbial Ecology in Premature Infants Assessed with Non-Culture-Based Techniques. J. Pediatr. 156, 20–25 (2010).

24. Keski-nisula, L. et al. Maternal intrapartum antibiotics and decreased vertical transmission of Lactobacillus to neonates during birth. Acta Peadiatrica 102, 480–485 (2013).

25. Fouhy, F. et al. High-throughput sequencing reveals the incomplete, short-term recovery of infant gut microbiota following parenteral antibiotic treatment with ampicillin and gentamicin. Antimicrob. Agents Chemother. 56, 5811–5820 (2012).

26. Gibson, M. K., Forsberg, K. J. & Dantas, G. Improved annotation of antibiotic resistance determinants reveals microbial resistomes cluster by ecology. ISME J. 9, 207–216 (2014).

27. Anderson, M. J. A new method for non-parametric multivariate analysis of variance. Austral Ecol. 26, 32–46 (2006).

28. Yatsunenko, T. et al. Human gut microbiome viewed across age and geography. Nature 486, 222–228 (2012).

29. Oh, J., Byrd, A. L., Park, M., Kong, H. H. & Segre, J. A. Temporal Stability of the Human Skin Microbiome. Cell 165, 854–866 (2016).

30. Schwarz, D. F., König, I. R. & Ziegler, A. On safari to Random Jungle?: a fast implementation of Random Forests for high-dimensional data. Bioinformatics 26, 1752–1758 (2010).

31. Strobl, C., Boulesteix, A.-L., Kneib, T., Augustin, T. & Zeileis, A. Conditional variable importance for random forests. BMC Bioinformatics 9, 307 (2008).

32. Szarecka, A., Lesnock, K. R., Ramirez-Mondragon, C. A., Nicholas, H. B. & Wymore, T. The Class D beta-lactamase family: residues governing the maintenance and diversity of function. Protein Eng. Des. Sel. 24, 801–809 (2011).

33. Wang, X. & Quinn, P. J. Progress in Lipid Research Lipopolysaccharide: Biosynthetic pathway and structure modification. Prog. Lipid Res. 49, 97–107 (2010).

34. Meredith, T. C., Aggarwal, P., Mamat, U., Lindner, B. & Woodard, R. W. Redefining the Requisite Lipopolysaccharide. ACS Chem. Biol. 1, 33–42 (2006).

35. Cech, D., Markin, K. & Ronald, W. Identification of a D-arabinose-5-phosphate isomerase in the Gram-positive Clostridium tetani. J. Bacteriol. (2017). doi:10.1128/JB.00246-17

36. Brown, C. T., Olm, M. R., Thomas, B. C. & Banfield, J. F. In situ replication rates for uncultivated bacteria in microbial communities. Nat. Biotechnol. 34, 57992 (2016).

37. Modi, S. R., Collins, J. J. & Relman, D. A. Antibiotics and the gut microbiota. J. Clin. Invest. 124, 4212–4218 (2014).

38. Piddock, L. J. V. Multidrug-resistance efflux pumps — not just for resistance. Nat. Rev. Microbiol. 4, 629–636 (2006).

39. Kanehisa, M. & Goto, S. KEGG: Kyoto Encyclopedia of Genes and Genomes. Nucleic Acids Res. 28, 27–30 (2000).

40. Zhu, J., Zou, H., Rosset, S. & Hastie, T. Multi-class AdaBoost. Stat. Interface 2, 349–360 (2009).

41. Valladares, M. H. et al. Zn(II) Dependence of the Aeromonas hydrophila AE036 Metallo-beta-lactamase Activity and Stability. Biochemistry 36, 11534–11541 (1997).

42. Hansen, L. H., Jensen, L. B., Sørensen, H. I. & Sørensen, S. J. Substrate specificity of the OqxAB multidrug resistance pump in Escherichia coli and selected enteric bacteria. J. Antimicrob. Chemother. 60, 145–147 (2007).

43. Jacoby, G. A. & Medeiros, A. A. More Extended-Spectrum Beta-Lactamases. Antimicrob. Agents Chemother. 35, 1697–1704 (1991).

44. Hughes, D. Exploiting genomics, genetics and chemistry to combat antibiotic resistance. Nat. Rev. Genet. 4, 432–441 (2003).

45. Arthur, M. & Quintiliani, R. Regulation of VanA-and VanB-Type Glycopeptide Resistance in Enterococci. Antimicrob. Agents Chemother. 45, 375–381 (2001).

46. Li, X.-Z. & Nikaido, H. Efflux-Mediated Drug Resistance in Bacteria: an Update. Drugs 69, 1555–1623 (2009).

47. DiGiulio, D. B. et al. Temporal and spatial variation of the human microbiota during pregnancy. Proc. Natl. Acad. Sci. 112, 11060–11065 (2015).

48. Yazdani, M. et al. Using Machine Learning to Identify Major Shifts in Human Gut Microbiome Protein Family Abundance in Disease. in IEEE International Conference on Big Data 1273–1280 (2016).

49. Sharon, I. et al. Time series community genomics analysis reveals rapid shifts in bacterial species, strains, and phage during infant gut colonization. Genome Res. 23, 111–120 (2013).

50. Kumar, V. et al. Comparative genomics of Klebsiella pneumoniae strains with different antibiotic resistance profiles. Antimicrob. Agents Chemother. 55, 4267–4276 (2011).

51. Kantor, R. S. et al. Bioreactor microbial ecosystems for thiocyanate and cyanide degradation unraveled with genome-resolved metagenomics. Environ. Microbiol. 17, 4929–4941 (2015).

52. Mousa, W. K. et al. Root-hair endophyte stacking in finger millet creates a physicochemical barrier to trap the fungal pathogen Fusarium graminearum. Nat. Microbiol. 1, 1–12 (2016).

53. Raveh-Sadka, T. et al. Gut bacteria are rarely shared by co-hospitalized premature infants, regardless of necrotizing enterocolitis development. Elife 2015, e05477 (2015).

54. Raveh-Sadka, T. et al. Evidence for persistent and shared bacterial strains against a background of largely unique gut colonization in hospitalized premature infants. ISME J. 10, 2817–2830 (2016).

55. Brooks, B. et al. Strain-resolved analysis of the hospital room and hospitalized infants reveals overlap between the human and room microbiome (in review).

56. Langmead, B. & Salzberg, S. L. Fast gapped-read alignment with Bowtie 2. Nat Methods 9, 357–359 (2012).

57. Peng, Y., Leung, H. C. M., Yiu, S. M. & Chin, F. Y. L. IDBA-UD: A de novo assembler for single-cell and metagenomic sequencing data with highly uneven depth. Bioinformatics 28, 1420–1428 (2012).

58. Hyatt, D. et al. Prodigal?: prokaryotic gene recognition and translation initiation site identification. BMC Bioinformatics 11, 119 (2010).

59. Zaharia, M. et al. Faster and More Accurate Sequence Alignment with SNAP. arXiv (2011).

60. Alneberg, J. et al. Binning metagenomic contigs by coverage and composition. Nat. Methods 11, 1144–1154 (2014).

61. Olm, M. R., Brown, C. T., Brooks, B. & Banfield, J. F. dRep: A tool for fast and accurate genome de-replication that enables tracking of microbial genotypes and improved genome recovery from metagenomes. ISME J. (2017).

62. Laing, C. et al. Pan-genome sequence analysis using Panseq?: an online tool for the rapid analysis of core and accessory genomic regions. BMC Bioinformatics 11, 461 (2010).

63. Finn, R. D., Clements, J. & Eddy, S. R. HMMER web server?: interactive sequence similarity searching. Nucleic Acids Res. 39, 29–37 (2011).

64. McArdle, B. & Anderson, M. Fitting Multivariate Models to Community Data?: A Comment on Distance-Based Redundancy Analysis. Ecology 82, 290–297 (2013).

65. Dixon, P. Computer program review VEGAN, a package of R functions for community ecology. J. Veg. Sci. 14, 927–930 (2003).

66. Pedregosa, F. et al. Scikit-learn: Machine Learning in Python. J. Mach. Learn. Res. 12, 2825–2830 (2012).

67. Edgar, R. C. MUSCLE: Multiple sequence alignment with high accuracy and high throughput. Nucleic Acids Res. 32, 1792–1797 (2004).

68. Stamatakis, A. RAxML version 8: a tool for phylogenetic analysis and post-analysis of large phylogenies. Bioinformatics 30, 1312–1313 (2014).

